# Model-based branching point detection in single-cell data by K-Branches clustering

**DOI:** 10.1101/094532

**Authors:** Nikolaos K. Chlis, F. Alexander Wolf, Fabian J. Theis

## Abstract

**Motivation:** The identification of heterogeneities in cell populations by utilizing single-cell technologies such as single-cell RNA-Seq, enables inference of cellular development and lineage trees. Several methods have been proposed for such inference from high-dimensional single-cell data. They typically assign each cell to a branch in a differentiation trajectory. However, they commonly assume specific geometries such as tree-like developmental hierarchies and lack statistically sound methods to decide on the number of branching events.

**Results:** We present K-Branches, a solution to the above problem by locally fitting half-lines to single-cell data, introducing a clustering algorithm similar to K-Means. These halflines are proxies for branches in the differentiation trajectory of cells. We propose a modified version of the GAP statistic for model selection, in order to decide on the number of lines that best describe the data locally. In this manner, we identify the location and number of subgroups of cells that are associated with branching events and full differentiation, respectively. We evaluate the performance of our method on single-cell RNA-Seq data describing the differentiation of myeloid progenitors during hematopoiesis, single-cell qPCR data of mouse blastocyst development and artificial data.

**Availability:** An R implementation of K-Branches is freely available at https://github.com/theislab/kbranches

## I. INTRODUCTION

Recent advances in single-cell technologies have lead to the discovery and characterization of novel cell types in multicellular organisms. Studying diverse cell populations that differ in morphology and function can pinpoint distinct cell types in different stages of regulatory processes, such as cellular development. For example, single-cell methods have lead to new discoveries related to hematopoietic stem cells [1, 2], as well as the immune system [3–5].

The development of novel computational techniques for the analysis of single-cell data is an active research topic in the field of bioinformatics [6–8]. The key idea is that individual cells can be mapped from a high dimensional space to a low-dimensional manifold of trajectories that reflect the continuous regulatory processes. As a result, a number of methods have been proposed that can reconstruct differentiation trajectories, given snapshot data of individual cells in different stages of the differentiation process, such as Monocle [9], Wishbone [10], Diffusion Pseudotime [11] and SLICER [12]. Given a “root” cell as a starting point, most of these methods can also calculate an ordering of the cells (pseudotime) based on the stage each cell is in the differentiation process. However, with the exception of Diffusion Pseudotime, while these methods are successful in assigning cells to discrete differentiation trajectories (branches) they do not tackle the problem of identifying the local dimensionality around each cell. That is, identifying branching regions of cells not yet strongly associated to any branch, intermediate regions along a branch and tip regions of fully differentiated cells. Moreover, all the above methods lack a sound statistical model to identify the existence and number of cell subgroups associated to branching events.

In this study, we propose a data driven, model-based clustering method that identifies the exact number of ’’branching regions”, as well as the exact number of fully differentiated “tip regions” in the lineage tree. The method then proceeds to assign each cell to a branching, intermediate or tip region. The proposed methodology does not aim to infer a pseudotemporal ordering of the cells and as such no “root” cell needs to be defined. Moreover, since characterization of each cell is based on local information in the differentiation trajectory, the method can successfully identify cells belonging to the aforementioned regions of interest in trajectories of arbitrary geometry.

## II. METHODS

### A. Problem formulation

Given a center **c** and direction **v**, a halfline *L* is defined as the set of points satisfying *L* = { **c** + *t*.**v**, *t* ≥ 0}, with **l**, **c**, **v** ∊ ℝ*^P^*. We aim to find *K* halflines *L_1_*, …, *L_K_* with a common center **c** and *K* distinct direction vectors **v**_1_, …, **v**_*K*_. In this case, each halfline *L_k_* corresponds one cluster *C_k_*. As a prerequisite to defining a cost function, note that the Euclidean distance of a given point **x** to a halfline *L_k_* reads:

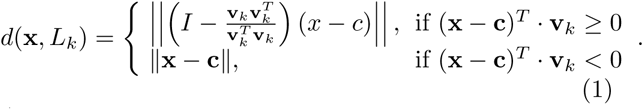

Additionally, one may also use other distance metrics[13].

The clustering method aims to assign each of the given data points (cells) into its closest halfline, while minimizing the total cost. In other words, the goal is to identify the center **c**, as well as the direction vectors **v**_1_, …, **v**_*K*_ of unit length that minimize the overall clustering cost. To this end, we define the cost function *J* to describe the total dispersion, which corresponds to the sum of dispersions over the *K* clusters and reads:

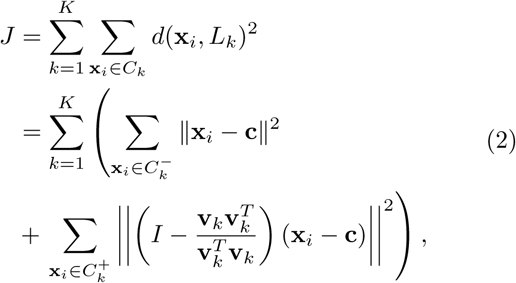

where 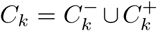 corresponds to all elements in cluster *k* and 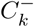, 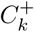 correspond to the sets of elements in cluster *k* with negative and positive dot product to to all vectors in the direction of *L_k_*, respectively.

**Algorithm 1.**
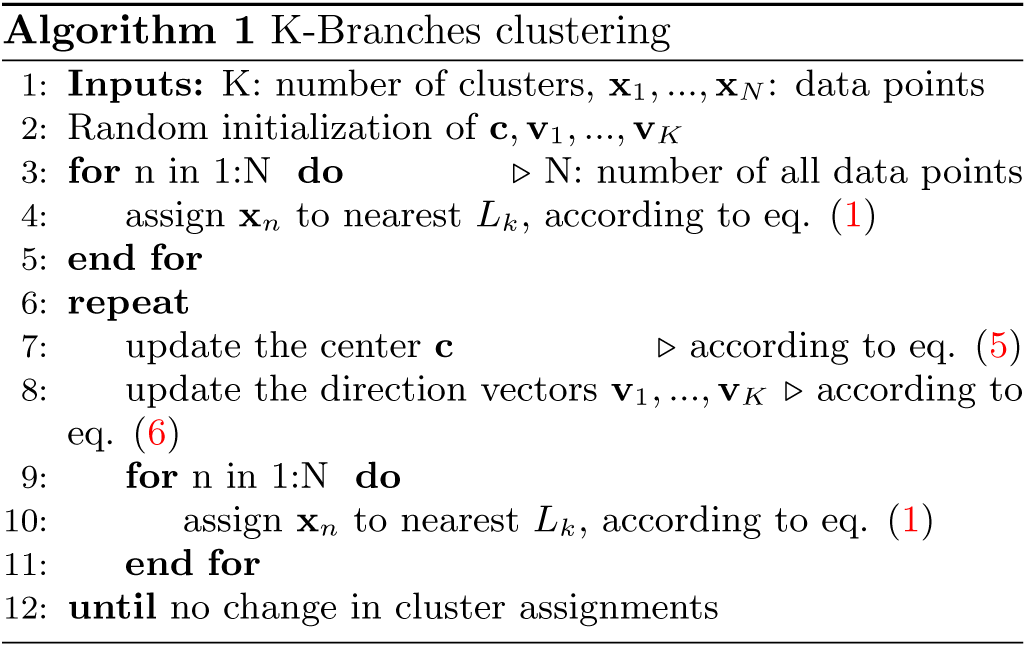
K-Branches clustering.

**Algorithm 2.**
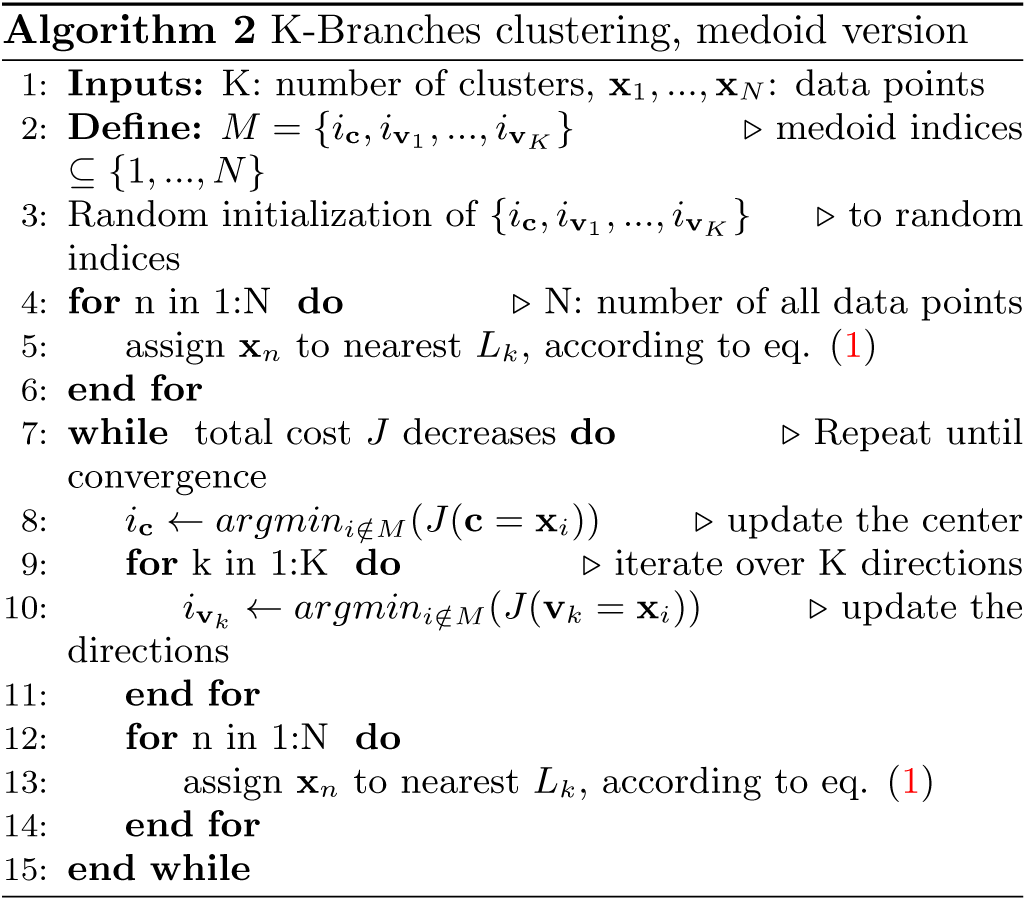
K-Branches clustering, medoid version.

#### 1. *The K-Branches clustering method*

In order to calculate the model parameters, after random initialization we follow an EM-like iterative optimization procedure similar to that of K-Means [14]. Namely, we iteratively (1) assign data points to their closest cluster and (2) update the estimates of **c** and **v**_1_, …, **v**_*K*_ while minimizing *J* in each step, until convergence. Since the method might converge to a local optimum of the cost function, multiple executions using different initializations have to be carried out. The method is randomly initialized by assigning one random data point as the center **c** and *K* other random data points as the direction vectors **v**_1_, …, **v**_*K*_. In the following subsections we present the update equations for the center and directions, respectively.

#### 2. *Estimating the center of the halines*

First, we optimize the cost function *J* with respect to the center of the halflines **c**. Therefore, we have to calculate the gradient ∇_**c**_ *J*, as follows:

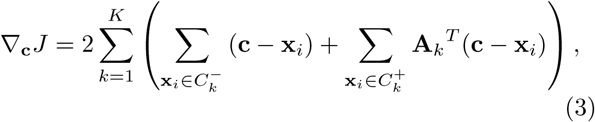

where the matrix **A_k_** is defined as:

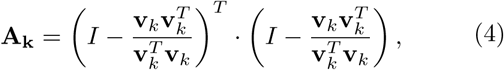

with 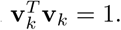

The equation ∇_c_*J* = 0 can be solved in closed form, and the optimal **c** reads:

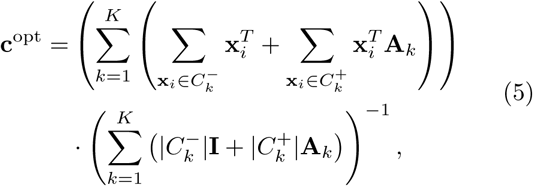

where 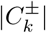 refers to the size of the set 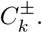 In the case *K* = 1 the right part of Equation (5) simplifies to 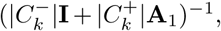 which is not full rank and therefore not invertible when 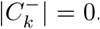 While the method for local clustering introduced in a subsequent section is also performed with *K* = 1, it uses a fixed center **c**, rendering the above limitation irrelevant.

#### 3. *Estimating the directions of the halflines*

The direction vector **v**_*k*_ for each of the K halflines (clusters) is updated according to Equation (6) as the average direction of all samples belonging to cluster *k*, with respect to the center of all halflines **c** and can subsequently be normalized to unit length.

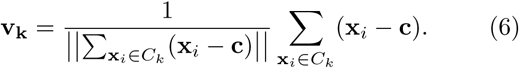

The pseudocode for the K-Branches algorithm is presented in **Algorithm 1**, while a comparison between K- Branches and K-Means is illustrated in Figure 1.

**FIG. 1.**
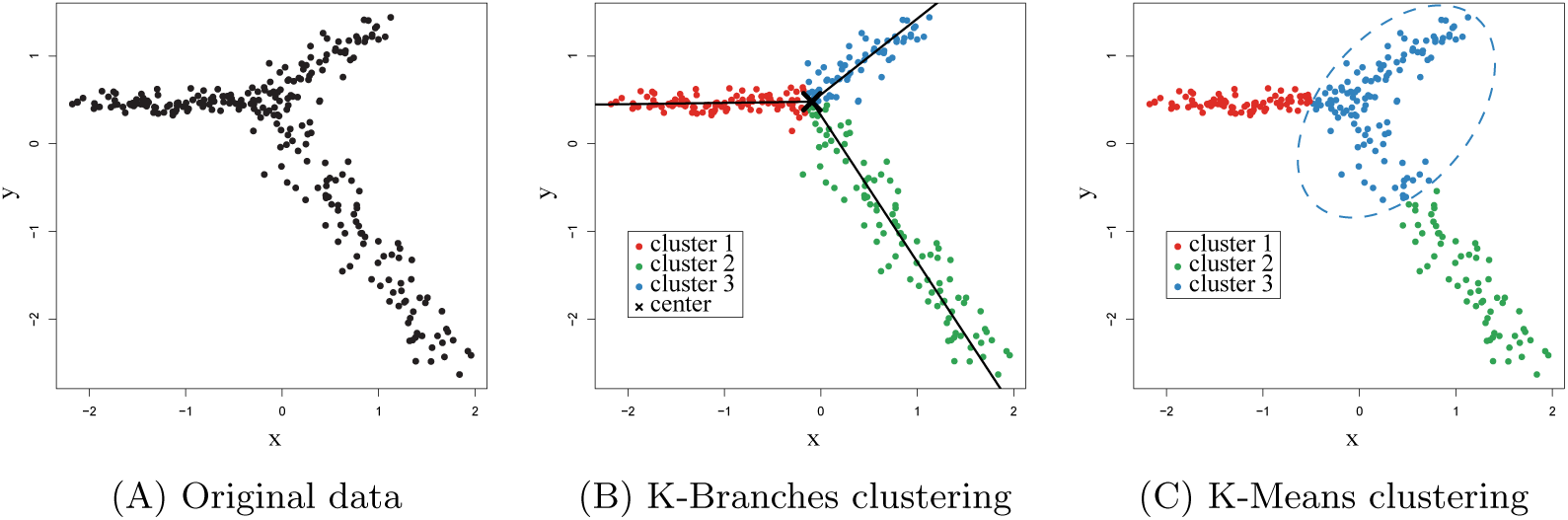
Illustration of K-Branches clustering on artificial data and comparison to K-Means. (A) Original data (B) In the case artificial data, K-Branches successfully clusters the three halflines. The center of the halflines as well as the lines corresponding to the direction of each cluster are plotted on top of the data points. The medoids version yields almost identical results for the same data. (C) Unlike K-Branches, K-Means (also with *K* = 3) merges part of the green halfline into the blue cluster. Since K-Means clusters points in spherical clusters, it is clearly not suitable for clustering data points which belong to distinct differentiation trajectories.

#### 4. *Medoid version of K-Branches*

As in K-Means, the K-Branches method described above determines a “centroid” *L_k_*(**c**, **v_k_**) per cluster, which depends on arbitrary vectors **c**, **v_k_** ∊ ℝ^*P*^. We can easily modify this to use data points, as in K-Medoids ([15]; [14]). The goal of the *Medoid* version of K-Branches is to identify one data point as the center *medoid* **x_c_** and *K* data points as the direction *medoids* **x**_**v**1_, …, **x**_**v***K*_. That is, the model parameters now correspond to *K* + 1 data points, instead of *K* +1 points in ℝ^*P*^, where *P* the number of dimensions. Similar to K-Medoids, the proposed algorithm searches over all data points during each iteration in a greedy manner and identifies the data points that minimize the cost function *J* given by Equation (2). All medoids are reassigned during each iteration of the algorithm, until a local minimum for *J* is reached and the total clustering cost cannot be further decreased. At this point, the algorithm converges to a solution where one of the data points is the center medoid **x**_c_ of the halflines and *K* data points correspond to the direction medoids **x_v_**_1_, …, **x**_**v***K*_. The relationship between the original and the medoid version is similar to that of K-Means and K-Medoids. That is, the medoid version is more robust in selecting the center of the halflines with respect to non-global optima and usually even only one random initialization is sufficient in practice. In the original algorithm, calculating the parameters **c**, **v**_1_, …, **v**_*K*_ requires time proportional to the number of data points *O*(*N*). A speedup of the medoid version is possible by computing the distance matrix **D** only once, where **D***_ij_* = ‖**x***_i_* – **x***_j_* ‖. Then, the distance of a data point **x***_i_* to a halfline *L*(**x_c_**, **x_v_** – **x_c_**) can be computed in O(1) time from Equation (7). However, for every one of the *N* – (*K* + 1) *candidate* medoids, the distance to every other data point is taken into consideration to calculate the overall clustering cost. As a result, *O*(*N*^2^) time is required to update the medoids during every iteration.

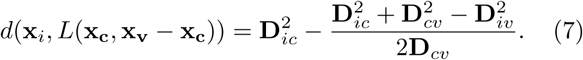

To summarize, in cases where robustness in the identification of the center of the halflines is crucial, the medoid version might be preferable. In applications where robust identification of the center of the halflines is not as crucial, especially in larger datasets, the original version of the algorithm could be preferable. Last, in cases where the center of the halflines is known (or held fixed), such as the case of local clustering presented later in the methods section, there is no advantage to using the medoid over the original version, since both are equally robust in identifying the directions of the halflines.

### B. Identifying branching and tip regions through local clustering

#### 1. *Local clustering*

In this section we derive a method for the identification of ’’regions of interest” in single-cell data, in particular, the identification of branching regions and tips of branches in lineage trees of differentiating single cells. The main idea is to center the previous model on each data point and adopt a local perspective by examining only the neighbourhood of *S* nearest neighbours to the center. We will show that by fixing the center of the halflines on a given data point and fitting the previous model of *K* halflines using a neighbourhood size of *S* data points, one can infer whether the center data point itself belongs to branching, intermediate or tip region, through appropriate model selection.

#### 2. *Selection of the neighbourhood size S*

The proposed method utilizes a number of *S* nearest neighbours to extract the neighbourhood of the cen- ter data point that is being examined. The size of the neighbourhood must be sufficiently large to reflect the local structure of the data, without capturing irrelevant global information. The proposed method is able to automatically suggest a value for *S* using a threshold on 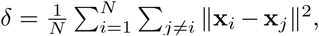 which ensures that the average cumulative squared distance *δ* of each data point to all other data points in the dataset is kept at a constant value. Moreover, the accompanying software package provides the option of visualization and manual fine tuning of *S* through a graphical user interface.

#### 3. *Neighbourhood scaling*

Another challenging aspect is related to datasets showing strong variation in the density of data points along the differentiation trajectories. For example, in the dataset of [16], there are sparse and dense regions. Variability of data point density might reflect an artifact of the data acquisition process, or could be a result of the underlying biological system. In the datasets examined so far, regions of very low density do not pose a threat to the performance of the method, since efficient selection of *S* will expand the neighbourhood size accordingly. On the other hand, the fixed number of *S* neighbours may drastically shrink the size of the neighbourhood in regions of very high density. To compensate for this effect, an appropriate heuristic rule was implemented. To be precise, for a given number of *S* neighbours, we calculate the median neighbourhood radius *ρ̅* over all neighbourhoods of size *S*. The neighbourhood scaling scheme is as follows: prior to performing local clustering for the *i*-th data point, its neighbourhood radius *ρ* (which corresponds to its distance to the furthest point in the neighbourhood) is calculated and the condition *ρ* ≥ *ρ̅* is assessed. If it is true, clustering is performed as usual. Otherwise, the neighbourhood size (*S*) of the i-th data point is increased until *ρ* ≥ *ρ̅* holds.

#### 4. *Local model selection*

The goal is to infer whether each data point belongs to a tip, intermediate or branching region of a differentiation trajectory, based on local clustering. That is, using a given data point as the fixed center **c** of the halflines, three different models are fit using *K*= 1, 2 and 3 halflines. The aim of the model selection step in the problem at hand is to identify the clustering model, i.e. the value of *K*, that best fits the data of the local neighbourhood centered around the data point in question. If one halfline best fits the neighbourhood, then the central data point belongs to a branch tip. If two halflines provide the best fit, then the central data point belongs to an intermediate region. If three halflines best fit the local neighbourhood, then the central data point belongs to a branching region. The concept of identifying regions of interest through model selection and local clustering is presented in Figure 2.

**FIG. 2.**
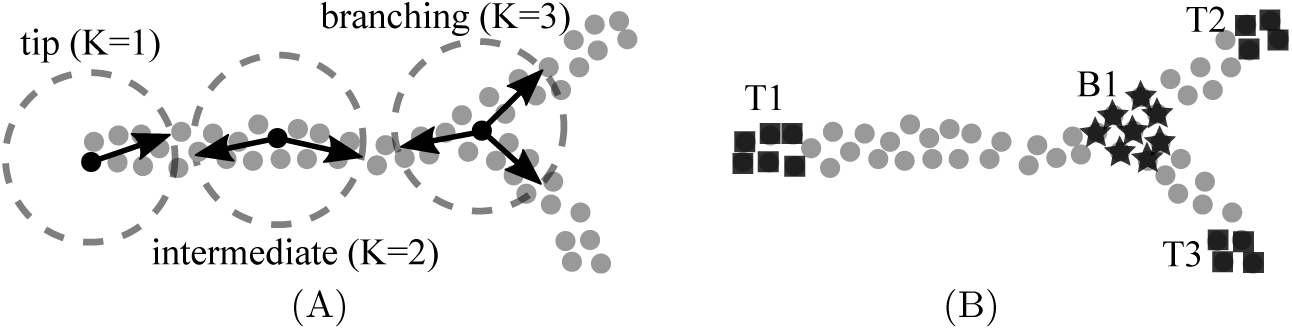
Illustration of local clustering for region identification. (A) Each cell is used as the center of the halflines and local clustering is performed in its neighbourhood. Then, by using model selection the center cell is either characterized as a tip cell, a cell belonging to an intermediate region or a cell belonging to a branching region depending on which of the three models (*K* = 1, 2, or 3) best describes the structure of the neighbourhood. (B) After local clustering is performed on the dataset, cells belonging to three tips (T1, T2, T3) and one branching region (B1) have been identified, while the rest of the cells are considered to belong to intermediate regions. The exact number of tip and branching regions is inferred from the data and does not need to be specified by the user.

The GAP statistic [17] is a popular method for identifying the number of clusters that best fit some given data. It depends on the sum of pairwise distances of points in each cluster. If the Euclidean distance is used as the distance measure, it corresponds to the dispersion around the cluster means (clustering cost). The GAP statistic compares the decrease in the clustering cost of the original data with the decrease in clustering cost of data drawn from a null distribution where no natural cluster structure exists. In theory, the dispersion in the data sampled from the null distribution decreases monotonically as *K* increases, while the dispersion in the original data drops rapidly for the value of *K* that best fits the dataset. Thus, the GAP statistic is maximized when the best value of *K* is used for clustering.

In the case of local K-Branches clustering, we introduce a modification of the GAP statistic that calculates the dispersion around halflines, as follows:

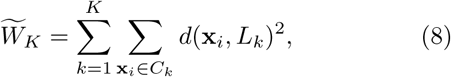

where *d*(***x***_*n*_, *L_k_*) is given by equation (1). Moreover, in contrast to the original GAP we do not take the logarithm of the dispersion, since it has been reported to overestimate the number of clusters in some cases [18]. Finally, the modified GAP statistic is given by:

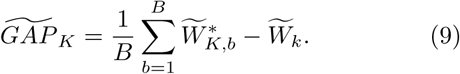

The dispersions 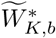 are calculated by applying eq. (8) after performing clustering on each of the *b* = 1, …, *B* bootstrap datasets (of the same size as the original dataset) drawn from the null reference distribution.

To summarize, given a data point as the center of the halflines, local clustering is performed. Then, if *GAP*_*K*=1_ > *GAP*_*K*=3_, it belongs to a branch tip. Otherwise, if the data point does not belong to a tip and 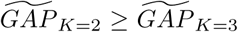 holds, it belongs to an intermediate region. Finally, if the data point does not belong to a tip and 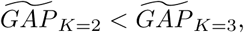 it belongs to a branching region. Both the original and modified versions of the GAP statistic are necessary for model selection and are complementary to each other. That is, *GAP* can identify tip cells but is not suitable for separating intermediate from branching cells (Figure 3-E). On the other hand, 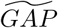 can separate intermediate and branching cells, but it not suitable for identifying tip cells, since it would falsely identify a large number of tip cells as branching cells (Figure 3-F). The performance comparison of the different GAP statistics is illustrated in Figure 3. As a final step, after all data points have been assigned to tip, intermediate and branching regions, K-Means clustering is performed on the subset of the data belonging to tips, using the original GAP statistic for model selection. In this manner, the exact number of tips is identified and each data point that has been characterized as belonging to a tip region is uniquely assigned to a specific tip. The same process is applied to cells belonging in branching regions in order to identify the exact number of branching events and assign ’’branching region” cells to their corresponding branching event.

**FIG. 3.**
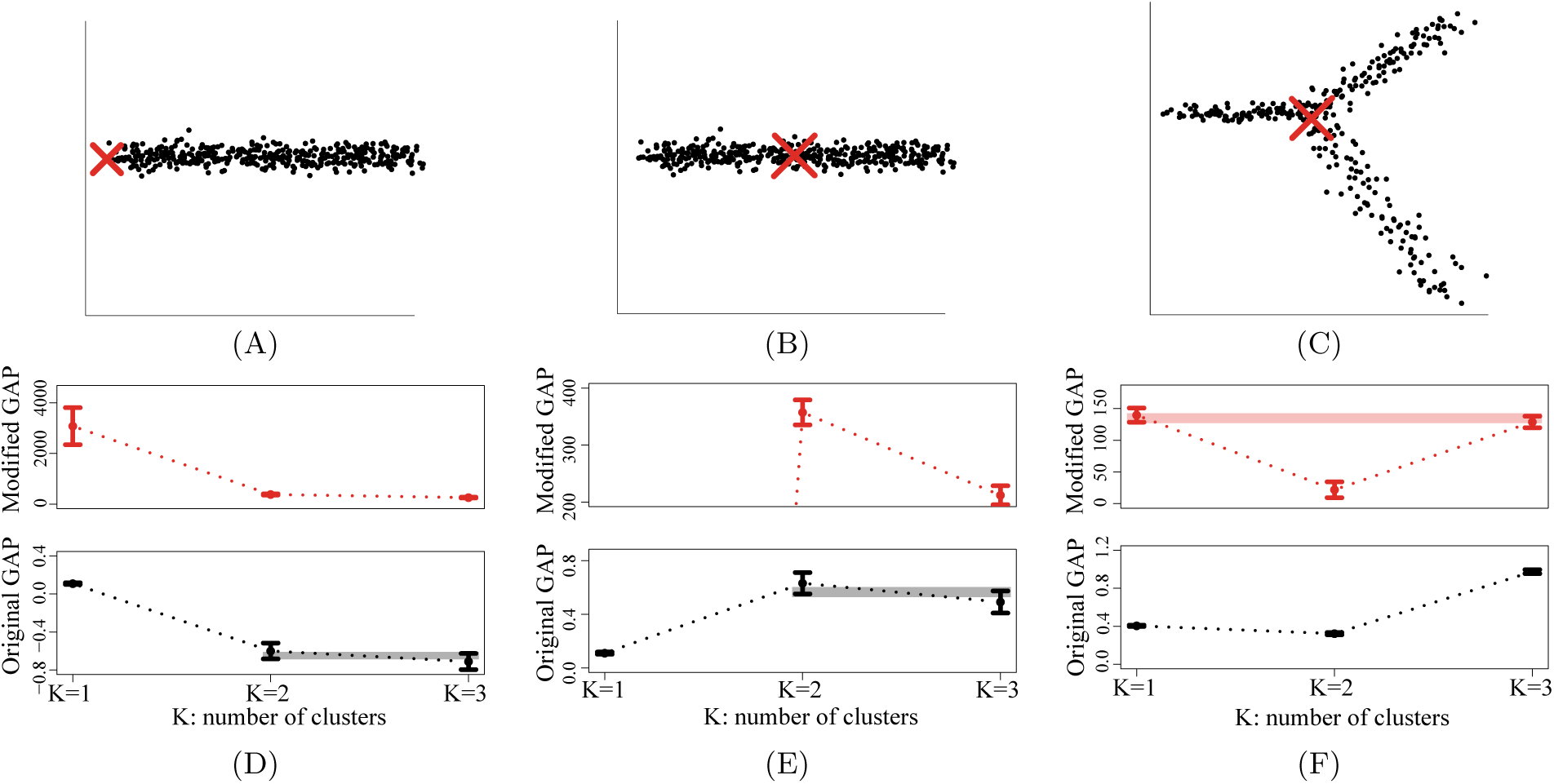
The original, as well as the modified versions of the GAP statistic are necessary for the identification of regions ofinterest in single-cell differentiation trajectories. All error bars in the plot correspond to 95% confidence intervals generatedusing 100 bootstrap data sets and overlaps of GAP statistics between different values of *K* are highlighted. Different y-axes areused since unlike the modified version, the original GAP statistic is calculated in log scale. (A) A local neighbourhood whichbelongs to a branch tip, where the center of the halines is fixed at the edge (red X) and its corresponding GAP statisticsin (D). (B) A local neighbourhood corresponding to an intermediate region is modelled by the same dataset, by moving the fixed center in the middle (red X) and its corresponding GAP statistics in (E): the overlap for GAP between *K* = 2 and *K* = 3 indicates it would lead to false identification of intermediate cells as branching cells (note: values of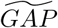 for *K* = 1 are very low and do not overlap with values for *K* = 2, 3). (C) Dataset showing a branching region and it’s corresponding GAPstatistics in (F): the overlap for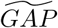 between *K* = 2 and *K* = 3 indicates it would lead to false identification of tip cells asbranching cells. These results indicate that GAP should be used first, to separate tip region cells from the rest (intermediateand branching). Subsequently, 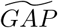 should be used to separate intermediate from branchign region cells.

#### 5. *Dimension reduction precedes model selection*

In this section we focus on the selection of the null reference distribution. Uniform sampling of features over a box aligned with the principal components of the data is suggested in [17]. Alternatively, uniform sampling over the range of every feature in the original dimensions of the data is suggested for simplicity. While the K-Branches clustering method performs well in the original space, model selection does not. This follows from the “curse of dimensionality” [14], since it becomes exponentially hard to estimate the null distribution in high dimensions. As a result, dimensionality reduction is a necessity if model selection is to be performed. Diffusion maps [19] are a non-linear dimensionality reduction method which are known to successfully identify differentiation trajectories [20], outperforming traditional dimensionality reduction methods such as principal component analysis (PCA) [14]. As a result, the data set is first processed by diffusion maps and the first few diffusion components (DCs) are selected. Then, local clustering is performed for each data point in the space of the selected DCs. Finally, the reference distribution is calculated by uniform sampling over a box aligned with the same DCs, resulting in the computation of the *GAP* and 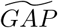statistics used for model selection.

## III. RESULTS

### A. Datasets

The proposed method is evaluated using two publicly available datasets, as well as one synthetic dataset. The first dataset corresponds to single-cell RNA-seq data describing the differentiation of myeloid progenitors during hematopoiesis ([2]; Accession Number GSE72857) and consists of measurements of 2730 cells and 8716 genes. The second dataset consists of single-cell qPCR data related to mouse blastocyst development ([16]; Accession Number J:140465) and includes measurements of 429 cells and 48 genes. Finally, the third dataset is an artificial dataset used as proof of concept and includes measurements of 2 synthetic genes and 244 cells that differentiate into three branches but the differentiation process includes a loop. Such a dataset could for example correspond to cellular reprogramming, or cells exiting the cell cycle, as also suggested by [12].

### B. Comparison to other methods

The purpose of local K-Branches is to identify branching and tip regions, while current popular methods assign cells to distinct branches. Local K-Branches is compared to Diffusion Pseudotime (DPT) [11] which in addition to assigning cells to distinct branches, also identifies tip cells and undecided cells in branching regions. One difference between DPT and the proposed method is that DPT only identifies one cell of each branch as the tip, while the proposed method typically identifies a region of tip cells. Monocle [9] and SLICER [12] are also indirectly compared to the proposed method, in terms of estimating correct branching in the data. The results of applying all methods on the above datasets are presented in Figure 4.

**FIG. 4.**
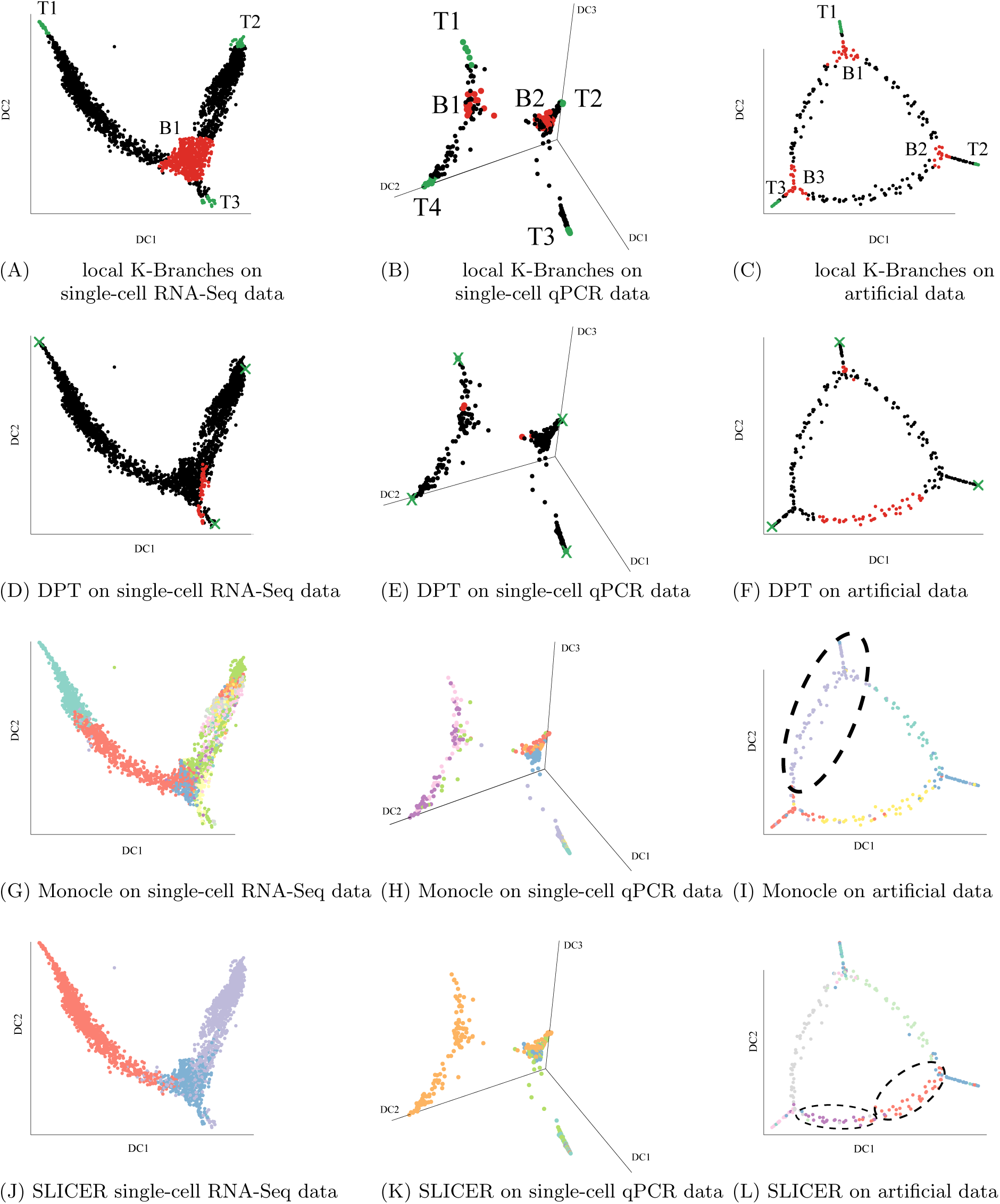
Local K-Branches successfully identifies regions of interest in single-cell data. Green, black and red points indicate cells belonging to tip (marked as T), intermediate and branching (marked as B) regions, respectively. In (G-L) the same diffusion components are used for visualization purposes only. (A) Local K-Branches identifies three tip regions and one branching region in single-cell RNA-Seq data [2]. (D) Diffusion Pseudotime yields similar results when applied on the same data. (B) Local K-Branches identifies four branch tips and two branching regions corresponding to two separate branching events in single-cell qPCR data [16]. (E) Once again, the results of Diffusion Pseudotime support the regions identified by K-Branches on the same data. (C) K-Branches successfully identifies three branch tips and three branching regions in artificial data, despite the loop between the branches. (F) Diffusion Pseudotime identifies the branch tips but fails to correctly identify all branching regions on the same data, due to the presence of the loop. (G-I) Monocle branch assignments on the same datasets. In (G, H) Monocle overestimates the number of branching events, while it does not detect one branching event in (I). (J-L) SLICER branch assignments on the same datasets. SLICER underestimates branching in (J), finds irrelevant branches in (K) and overestimates branching in (L).

The comparison to a DPT analysis was performed as follows: First, DPT was applied to each dataset and the cells corresponding to the tips of the branches, as well as the undecided cells corresponding to branching regions were identified. To ensure that the comparison is as direct as possible the same diffusion components computed by diffusion map during DPT analysis were extracted. Subsequently, local K-Branches clustering was performed on exactly the same diffusion components and cells were assigned to either fully differentiated tip, intermediate and branching regions. It should be noted that while DPT uses all available diffusion components, local K-Branches is only performed on the first two or three components, depending on the morphology of the dataset.

#### 1. *Single-cell RNA-Seq data of myeloid progenitors*

When applied to the first two diffusion components of the single-cell RNA-Seq dataset of [2], the proposed method identifies three branch tips of fully differentiated cells, as well as one branching region. The results of Diffusion Pseudotime on the same data agree with the findings of the proposed method. Two of the three tips identified by DPT are in the tip regions of local K-Branches, while the third tip of DPT is not inside but in the vicinity of the local K-Branches tip region. When comparing the branching region, the undecided cells of DPT are either inside or in close proximity to the branching region identified by local K-Branches. However, considerably fewer cells are considered as undecided by DPT. Finally, Monocle overestimates, while SLICER underestimates the overall branching. The regions identified by K-Branches are illustrated with respect to FACS labels in Figure 5.

**FIG. 5.**
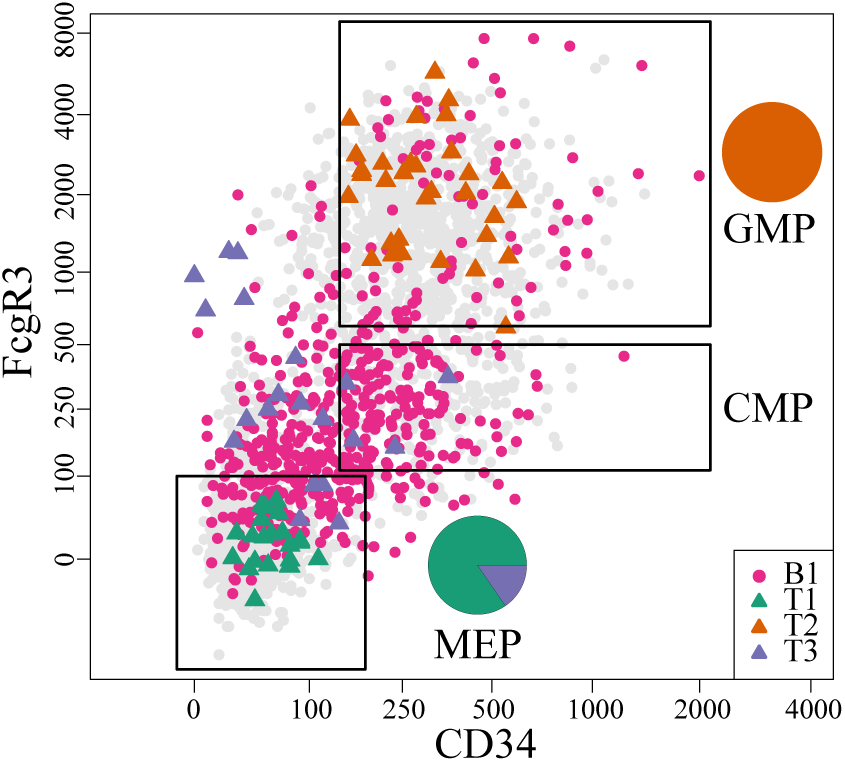
Cells plotted according to FACS-measured FcgR3 and CD34 protein expression values [2]. The cells corresponding to regions B1, T1, T2, T3, as identified by local K- Branches, are highlighted. The megakaryocyte/erythrocyte progenitors (MEP), granulocyte/macrophage progenitors (GMP) and common myeloid progenitors (CMP) gates are also plotted. Pie-charts correspond to the distribution of T1,T2 and T3 cells in the MEP and GMP gates. The cells of branching region B1 are enriched only in the CMP gate (Fisher’s exact test p-value: 2.2^−16^). The cells of tip T1 correspond to MEP, while the cells of tip T2 correspond to GMP. The cells of tip T3 correspond to outlier groups of dendritic cells and natural killer cells (lymphocytes).

#### 2. *Single-cell qPCR data of mouse blastocyst development*

The proposed method was applied to the first three diffusion components of the single-cell qPCR data which contains two distinct branching events [16]. Once more there is close agreement between the results of the proposed method and Diffusion Pseudotime. Both methods identify four branch tips and the tip cells of DPT are in the tip regions of the proposed method. Moreover, both methodologies identify two branching regions indicating two separate branching events. However, DPT only assigns two cells per branching region, while the branching regions identified by the proposed methodology are considerably larger. Finally, one key difference is that the proposed method automatically identified four branch tips and two branching regions, while DPT had to be manually run twice on the data: First, three branches were identified, then DPT was performed on one of the branches, identifying the second branching region and new branch tips. Both Monocle and SLICER do not identify the two branching events, probably due to the high degree of sparsity in the data.

#### 3. *Artificial data of arbitrary geometry*

The third dataset highlights an important advantage of the proposed methodology. Namely, the identification branch tips and branching regions in datasets of arbitrary geometry. In this case, the dataset was manually generated to consist of three branches with a loop among them and the first two diffusion components retain the same geometry as the original dataset. Even though it could be directly applied to the original two-dimensional data, the proposed method was performed on the first to diffusion components. This was done for two reasons: First, for real data of high dimensions clustering and model selection will be performed on the diffusion components and we assume that dimensionality reduction through diffusion maps will also retain the loop structure of real data. Second, by using the diffusion components there is direct comparison to the performance of DPT. Despite the challenging geometry of the dataset, the proposed method correctly identifies the three regions corresponding to the branch tips, as well as the three branching regions. On the other hand, DPT correctly identifies the three tip cells but fails in identifying the branching regions. To be precise, it identifies one branching region correctly, but then it fails to find the other two and considers one irrelevant part of the loop as a branching region. Monocle underestimates the number of branching events, probably since it always assumes that the differentiation trajectory corresponds to a tree-like structure. Finally, SLICER overestimates the overall branching in the data.

## IV. CONCLUSION AND DISCUSSION

In this study, a model based clustering approach was introduced for the identification of regions of interest in single-cell data. First, a novel clustering method called K-Branches was introduced, which clusters data points into a set of K halflines with a common center. Subsequently, this clustering method was applied locally to the neighbourhood of each cell and a modified version of the GAP statistic was developed to perform model selection. The goal of model selection is to identify the *local dimensionality* of the data. That is, identify fully differentiated tip cells and cells belonging to branching regions. In this manner, all branching events, as well as all end-points (tips) in differentiation trajectories can be identified. As demonstrated, this local view of the data allows the method to be successfully applied to challenging datasets that include sparsity and complex geometries.

The main idea of the proposed methodology is different from that of commonly used methods such as DPT, Monocle, Wishbone, or SLICER. To be precise, these methods aim to assign each cell to a distinct branch in the differentiation process and also calculate pseudotime: an ordering of the cells, relevant to their distance from a starting root cell, which reflects how far they have progressed in the differentiation process. As such, K-Branches cannot be directly compared to most of these methods, perhaps with the exception of DPT. To be precise, DPT also identifies tip cells and branching regions of undecided cells.

The performance of the proposed method was compared to that of DPT in three single-cell datasets. In the fist two datasets which correspond to single-cell qPCR [16] and single-cell RNA-Seq [2] data, both methods yield similar results. The main differences being that DPT assigns fewer cells in branching regions and only assigns one cell per tip. However, in the third dataset which consists of three branches with a loop in between, the model based approach of the proposed methodology successfully identifies all tip and branching regions, while DPT only identifies the branch tips and does not manage to correctly identify the branching regions of undecided cells. While this difference was observed on a synthetic data set, real datasets containing loops could in theory correspond to cells exiting cell cycle, cells resulting in the same state through different differentiation trajectories, or cellular reprogramming [21]. One advantage of DPT is faster execution time since the entire dataset is typically processed in a few minutes. On the other hand, local K-Branches requires a few seconds *per data point*. However, in the case of local K-Branches each data point can be processed completely in parallel.

In terms of future work, it would be interesting to extend the method to support explicit identification of the branches that lie between the branching and tip regions, which are currently only characterized as intermediate regions. While clustering works in the original dimensions, model selection using the GAP statistic does not. As such, the proposed method utilizes diffusion maps for dimensionality reduction. In this regard coupling the proposed methodology with dimensionality reduction methods other than diffusion maps and comparing the performance achieved would be interesting. Finally, developing a different model selection method, other than the GAP statistic, that would allow the methodology to be directly applied in the original dimensions could be an additional topic of future work.

## ACKNOWLEDGEMENTS

We would like to acknowledge L. Haghverdi for her helpful advice. We would like to thank M. Buttner for her comments and support on drawing biological conclusions. Finally, we would like to thank P. Angerer and D. S. Fischer for their comments on the R package and manuscript, respectively.

